# Improved genome assembly of American alligator genome reveals conserved architecture of estrogen signaling

**DOI:** 10.1101/067165

**Authors:** Edward S. Rice, Satomi Kohno, John St. John, Son Pham, Jonathan Howard, Liana Lareau, Brendan O’Connell, Glenn Hickey, Joel Armstrong, Alden Deran, Ian Fiddes, Roy N. Platt, Cathy Gresham, Fiona McCarthy, Colin Kern, David Haan, Tan Phan, Carl Schmidt, Jeremy Sanford, David A. Ray, Benedict Paten, Louis J. Guillette, Richard E. Green

## Abstract

The American alligator, *Alligator mississippiensis*, like all crocodilians, has temperature-dependent sex determination, in which the sex of an embryo is determined by the incubation temperature of the egg during a critical period of development. The lack of genetic differences between male and female alligators leaves open the question of how the genes responsible for sex determination and differentiation are regulated. One insight into this question comes from the fact that exposing an embryo incubated at male-producing temperature to estrogen causes it to develop ovaries. Because estrogen response elements are known to regulate genes over long distances, a contiguous genome assembly is crucial for predicting and understanding its impact.

We present an improved assembly of the American alligator genome, scaffolded with *in vitro* proximity ligation (Chicago) data. We use this assembly to scaffold two other crocodilian genomes based on synteny. We perform RNA sequencing of tissues from American alligator embryos to find genes that are differentially expressed between embryos incubated at male-versus female-producing temperature. Finally, we use the improved contiguity of our assembly along with the current model of CTCF-mediated chromatin looping to predict regions of the genome likely to contain estrogen-responsive genes. We find that these regions are significantly enriched for genes with female-biased expression in developing gonads after the critical period during which sex is determined by incubation temperature. We thus conclude that estrogen signaling is a major driver of female-biased gene expression in the post-temperature sensitive period gonads.

## Introduction

The American alligator, *Alligator mississippiensis*, like all crocodilians and many other reptiles, has temperature-dependent sex determination, in which the sex of an embryo is determined by the incubation temperature of its egg during a temperature-sensitive period (TSP) of development (Ferguson and Joanen 1982). In contrast, mammals, birds, and other animals with genetic sex determination have different combinations of sex chromosomes between the two sexes. These genetic differences between males and females determine the sex of an individual, and induce sex differentiation during development by causing differential expression of numerous genes. Genes with sex-biased expression during development in these lineages include conserved sexual development genes such as *SOX9* and *WNT4*. Such expression differences eventually cause the development of one of two sets of distinct sexual characteristics. However, in alligators and other species with temperature-dependent sex determination, males and females have identical genomes, leaving open the question of how differences in temperature lead to differential expression of genes between males and females during early development (Morrish and Sinclair 2002; Shoemaker-Daly et al. 2010; Kohno and Guillette 2013).

One insight into this question comes from the observation that exposing an alligator embryo to estrogen while incubated at a male-producing temperature causes it to develop ovaries instead of testes via the transcription factor estrogen receptor alpha (Bull et al. 1988; Milnes et al. 2005; Kohno et al. 2015), demonstrating that estrogen signaling is an early effector of sexual development genes in the American alligator. In addition, *CYP19A1*, the gene coding for the enzyme aromatase, which converts androgen to estrogen, is expressed at significantly higher levels in embryos incubated at female-producing temperature than those incubated at male-producing temperature (Gabriel et al. 2001). It remains unknown which genes estrogen regulates to induce ovarian development, or the mechanism by which it targets these genes via estrogen receptor alpha.

In humans, estrogen regulates gene expression through several mechanisms, most of which involve the transcription factors estrogen receptor alpha and beta. In the most well-known of these mechanisms, the estrogen 17β-estradiol activates an estrogen receptor by binding to its ligand-binding domain, thus allowing the receptor’s DNA-binding domain to bind to a well-defined enhancer sequence, the estrogen response element, promoting the expression of nearby genes (Nilsson et al. 2001; Dahlman-Wright et al. 2006). The motif to which human estrogen receptor alpha binds has been well-characterized using chromatin immunoprecipitation (Gruber et al. 2004; Laganière et al. 2005). A majority of estrogen receptor alpha binding sites are distal enhancers–that is, they are far from the genes they regulate (Carroll et al. 2006; Lin et al. 2007; Welboren et al. 2009). Further, a majority of estrogen receptor binding events are associated with long-range intrachromosomal chromatin interactions, and these associated events are significantly enriched for RNA polymerase II recruitment (Fullwood et al. 2009). The zinc-finger protein *CTCF* is responsible for many of these chromatin interactions (Zhang et al. 2010). Regions delineated by two *CTCF* binding, that contain an estrogen receptor binding site are significantly more likely to contain estrogen-responsive genes in human (Chan and Song 2008). Because the estrogen response is a long-range phenomenon in humans, it suggests that a contiguous genome assembly is necessary to fully explore the genome architecture of estrogen regulation in alligators.

Green et al. (2014) published the genomes of the American alligator and two other crocodilians: the saltwater crocodile *Crocodylus porosus* and the gharial *Gavialis gangeticus*, with scaffold N50s of 205 kb and 127kb, respectively. The slow rate of molecular evolution within crocodilians (Green et al. 2014) makes this clade an ideal one for testing the ability to use a highly-contiguous genome assembly to scaffold the genome assemblies of related organisms based on synteny.

We present an improved American alligator genome assembly with scaffold N50 increased from 508 kbp to over 13 Mbp using Chicago, a recently-published technique for scaffolding genomes using *in vitro* proximity ligation read pairs (Putnam et al. 2016). We use the improved alligator genome to scaffold two other crocodilian genomes by aligning their scaffold to the alligator genome to infer adjacent scaffolds. With gene expression data from RNA-sequencing of three embryonic alligator tissues incubated at either male-producing or female-producing temperature, we find that genomic regions predicted by our model to be domains of estrogen regulation of gene expression are significantly enriched for genes with female-biased expression in the post-temperature sensitive period embryonic gonads.

## Results

### Assembly and annotation

The updated American alligator genome assembly has a total length of 2.16 Gbp compared to 2.17 Gbp for the previously published assembly. However, the new genome shows a 25-fold improvement in scaffold N50, a measure of contiguity, from 508 kbp to over 13 Mbp.

To assess the quality and accuracy of the new assembly, we measured its concordance with previously-published BAC-end pairs (Shedlock et al. 2007) that were not used in the assembly or scaffolding. Using bwa mem (Li 2013) with default parameters, we aligned the forward and reverse reads of the 1309 BAC-end pairs to the new assembly and to the assembly prior to scaffolding using Chicago data. We found that while 142 BAC-end pairs had both ends aligning to the same scaffold of our assembly before scaffolding with Chicago, 1160 BAC-end pairs have both ends aligning to the Chicago-scaffolded assembly. 1143, or 98.5%, of these pairs aligning to the same scaffold are oriented correctly, and 1125, or 98.4%, of these correctly-oriented pairs have an insert size between 70kb and 180kb. We thus conclude that this assembly is both accurate and an improvement over assembly not using the Chicago library.

We annotated the genome for protein-coding genes using previously-published RNA-seq reads (Green et al. 2014) and AUGUSTUS (Stanke et al. 2006), finding 32,052 transcripts and 24,713 genes. Moreover, we were able to assign names to 15,977 of these genes based on orthology with named genes in other vertebrate species. Using both orthology and protein sequence analysis, we assigned 5,960 unique Gene Ontology (GO) terms to 17,430 American alligator proteins.

### Crocodilian versus mammalian genome synteny

While the previous assembly of the American alligator genome (Green et al. 2014) was sufficient to compare to other genomes at the sequence level, our new long-range assembly presents an opportunity to perform genome comparisons on a broader scale. We computed synteny between the alligator and chicken (Galgal4) genomes using SyMAP v4.2 (Soderlund et al. 2011). We used both the new alligator assembly and the previous assembly for comparison, and found that the increased contiguity of the new assembly vastly improved our ability to compute synteny between the chicken and alligator genomes, more than doubling the percentage of the genome covered by synteny blocks from 35% to 90% and increasing the sizes of synteny blocks, with 57 of the 90 synteny blocks over 10Mb in length. A dot plot of synteny over the entire genomes shows that most scaffolds in the new alligator assembly correspond to a contiguous region of a chicken chromosome, although often with some intra-chromosomal rearrangements (Figure 1a,b). Some scaffolds in the alligator genome appear to correspond to whole arms of chicken chromosomes. For example, two alligator scaffolds almost completely cover GGA7. Furthermore, the microchromosome GGA10 is almost fully covered by a single alligator scaffold, scaffold 10 (Figure 1d), with one large inversion and numerous small local inversions.

**Figure 1.**
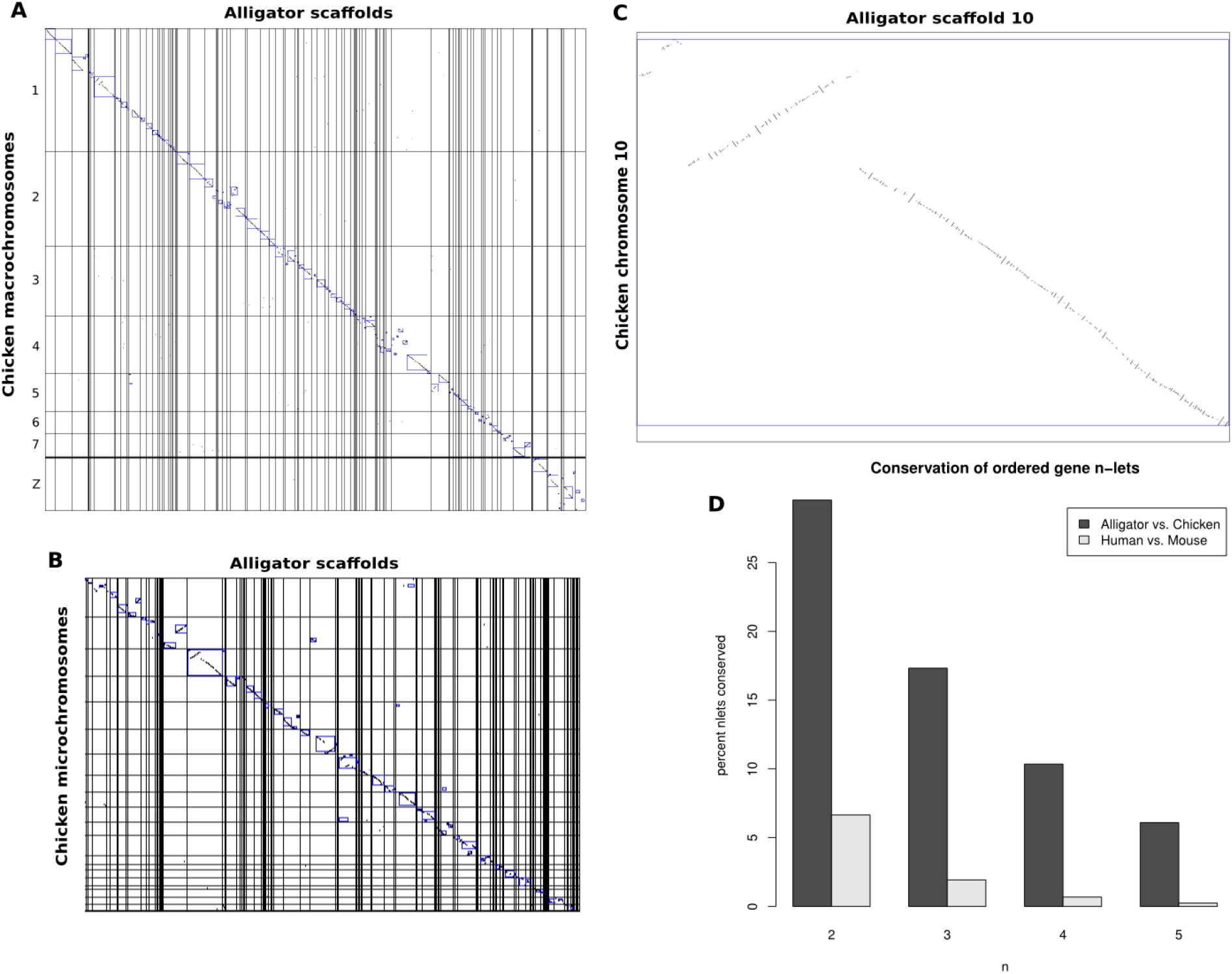
Our new long-range assembly of the American alligator genome allows analysis of the synteny between crocodilians and birds. (a-b) Dot plots of an anchored whole-genome alignment between the chicken and American alligator genomes shows a high degree of synteny, with many long alligator scaffolds covering significant portions of chicken chromosomes, including macrochromosomes (a) and microchromosomes (b). (c) Alligator scaffold 10 covers a vast majority of the chicken microchromosome 10. However, there are numerous small inversions and one large inversion between the two. (d) Conservation of ordered gene doublets, triplets, quadruplets, and quintuplets between alligators and chickens vs. between humans and mice, showing much higher synteny between alligators and chickens than between humans and mice.

To contrast the levels of genome rearrangement in archosaurs and mammals, we compared conservation of gene order between alligators and chickens (242 MYA TMRCA) to that between humans and mice (110 MYA TMRCA) (Crottini et al. 2012). We calculated the percentage of ordered pairs, triplets, quadruplets, and quintuplets of directly adjacent genes that occur in both alligators and chickens, and both humans and mice. We found four times greater conservation of gene pair synteny between alligators and chickens than between humans and mice, nine times greater conservation of gene triplets, fifteen times greater conservation of quadruplets, and twenty-five times greater conservation of quintuplets (Figure 1d).

A closer look at synteny between the chicken Z chromosome and the alligator genome reveals the expected inversion around the avian sex-determining gene *DMRT1* (Figure S4). This is concordant with the Z-linked inversions previously predicted by examining gene synteny between the avian Z chromosome and other reptilian outgroups such as the green anole *Anolis carolinensis*, red-tailed boa *Boa constrictor*, and Mexican musk turtle *Staurotypus triporcatus* (Kawagoshi et al. 2014; Zhou et al. 2014).

### Sex-biased gene expression

A crucial step towards understanding temperature-dependent sex determination in the American alligator is determining what genes are turned on or off based on temperature at various developmental stages. This necessitates the generation of a catalog of genes that show significantly different expression between eggs incubated at male-producing temperature and those incubated at female-producing temperature. To this end, we incubated alligator eggs at male- and female-producing temperatures for either zero, three, or thirty days after developmental stage 19. The period during which an embryo’s sex is determined by incubation temperature (temperature sensitive period; TSP) spans developmental stages 21 to 24 (Lang and Andrews 1994), which occur between our three- and thirty-day timepoints. We harvested the embryos after incubation, sub-dissected the gonad-adrenal-mesonephros complex into its constituent parts, and performed RNA sequencing on each of these three tissues for each sample. See Table S1 for a list of libraries sequenced along with their NCBI accessions.

We used the resulting RNA-seq data to quantify gene expression and determine which genes are differentially expressed between developing male and female embryos at these developmental stages in these three tissues. We found many genes with differential expression between males and females in each tissue at both the three-day and thirty-day timepoints (Figure S2). Unsurprisingly, the gonads at the post-TSP time point displayed the most sexual dimorphism in gene expression. The genes differentially expressed between male and female embryos in these samples include many genes known to be involved in early sexual development in other vertebrates. Such male development genes include *SOX9*, which triggers testis formation, and *AMH*, which inhibits the formation of Müllerian ducts (De Santa Barbara et al. 1998). Female development genes with female-biased expression in the post-TSP gonads include *CYP19A1*, which produces aromatase, the enzyme that converts androgens to estrogens (Toda and Shizuta 1993), and *FST*, which inhibits the production of follicle-stimulating hormone (Ying et al. 1987). *CYP19A1* was the gene with the most female-biased expression, with a log-2 fold-change of 12.463. This is consistent with other studies of aromatase expression in embryos incubated at different temperatures (Smith et al. 1995; Gabriel et al. 2001). *ESR1*, the gene coding for estrogen receptor alpha, and *CTCF* are highly expressed in both male and female gonads at this time point, with respective average FPKM values of 24.08 and 47.02, but no significant sex bias.

*TRPV4* has been suggested as one thermosensitive gene involved in TSD in the American alligator (Yatsu et al. 2015). We found no significant expression or sex-bias of *TRPV4* at any of our time points in any of the three tissues. This may be due to our sampling at different developmental stages than Yatsu et al. (2015).

### Estrogenic regulation of gene expression

Estrogen regulation of gene expression is best understood in humans from work dissecting the molecular basis of estrogen-responsive and non-responsive breast cancers in tissue models. That work has shown that in human estrogen-responsive tissues, estrogen promotes the expression of genes by allowing estrogen receptors to bind to enhancer DNA sequences (Dahlman-Wright et al. 2006). However, the enhancers to which estrogen receptors bind are usually distal to the genes they regulate (Carroll et al. 2006). Due to the sex-reversing effects of estrogen exposure during crocodilian development via estrogen receptor alpha (Kohno et al. 2015) and the extreme female-biased expression of the gene coding for aromatase, we hypothesize that estrogen signaling through *ESR1* binding is a major driver of female-biased gene expression during temperature-dependent sex determination in the American alligator.

We first tested this hypothesis by looking for enrichment of genes with female-biased expression in the post-TSP gonads of alligator embryos in the genomic regions surrounding computationally-predicted estrogen receptor binding sites. The DNA-binding domain of *ESR1* is perfectly conserved among humans, chickens, and alligators (Figure S3b, Table S2) and the DNA-binding motif of *ESR1* in human estrogen-responsive cells is well characterized (Gruber et al. 2004; Carroll et al. 2006; Lin et al. 2007). Therefore, we predicted *ESR1* binding sites in the American alligator genome using the motif representing the human estrogen response element. We found that while 337 (2.26%) of the 14,943 genes expressed in the post-TSP gonad have female-biased expression, 62 (3.11%) of the 1,991 expressed genes within 50kb of a putative estrogen receptor binding site have female-biased expression (E = 44.9; hypergeometric test p = 4.79 × 10^−3^). This indicates that genes are significantly more likely to have female-biased expression in the post-TSP gonad if they are near a location in the genome where *ESR1* is predicted to bind.

However, in human tissue models of estrogen regulation of gene expression, whether a gene is likely to be estrogen-responsive is based on its genomic location relative to not only estrogen receptor binding sites, but also *CTCF* binding sites (Chan and Song 2008). Since this was established in 2008, studies of CTCF-mediated chromatin looping have shown that *CTCF* helps divide the genome into functional domains through a chromatin extrusion process that causes loops to form only where two adjacent *CTCF* binding motifs are oriented towards each other (Rao et al. 2014; Sanborn et al. 2015). For a given pattern of *CTCF* binding motifs, there are often multiple possible looping topologies, and in practice, each of these occurs to some degree in a population of cells.

*CTCF* binding sites in the chicken genome are well-established (Martin et al. 2011) and the zinc finger domains of *CTCF* are perfectly conserved among human, chicken, and alligator orthologs (Figure S3a). We therefore used the *CTCF* binding motif in the chicken genome to predict *CTCF* binding sites in the American alligator genome. We used these binding site predictions and the most recent model of CTCF-mediated chromatin looping (Sanborn et al. 2015) to predict how chromatin loops form in the alligator genome (Figure 2a). We predicted 19,482 chromatin loops based on *CTCF* binding sites, comparable to the 21,306 found experimentally in the human genome (Li et al. 2012). 3,758 (19.3%) of these putative loops contain one or more *ESR1* binding sites. 10,074 (67.4%) of the 14,943 genes expressed in gonads after thirty days of incubation are within the boundaries of one or more predicted *CTCF* loops.

**Figure 2.**
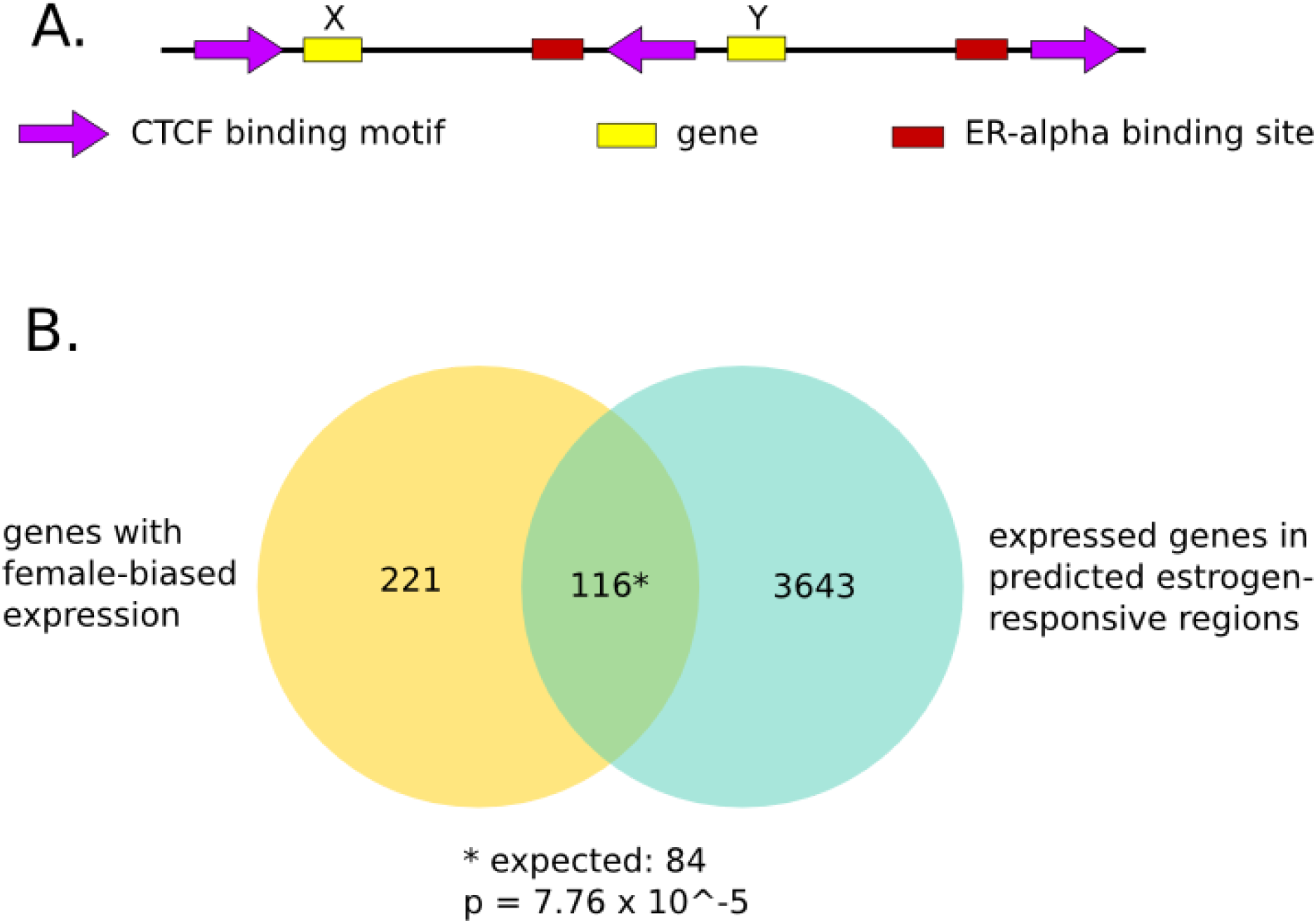
Genes in regions of the genome predicted to be under estrogenic regulation of gene expression are significantly more likely to be female-biased in the post-temperature sensitive period gonads. (a) Our model for predicting regions of the genome under estrogenic regulation of gene expression, based on the CTCF extrusion model (Sanborn et al. 2015) and the Chan & Song model of estrogen receptor binding site activity (Chan and Song 2008). In this example, Gene X is predicted to be estrogen-responsive and Gene Y is not because Gene X is between two inward-oriented CTCF binding motifs along with an ER binding site, while Gene Y is not. (b) Of the 14,943 genes expressed in the post-temperature sensitive period gonads, 337 have female-biased expression and 3,759 are in predicted estrogen-responsive genomic regions. However, 116 of these genes are both female-biased and within predicted estrogen-responsive regions, a significantly higher number than the expected 84 (p = 7.76 × 10^−5^).

We found that while 337 (2.26%) of the 14,943 genes expressed in the post-TSP gonads have female-biased expression, 116 (3.09%) of the 3759 expressed genes in CTCF loops containing one or more *ESR1* binding sites have female-biased expression (E = 84, hypergeometric test p = 7.76 × 10^−5^). This shows a significant enrichment in female-biased gene expression in the regions of the genome predicted to be estrogen-responsive under our model, providing support for our hypothesis that many of these genes are regulated by estrogen during sexual differentiation and development (Figure 2b).

Among the female-biased genes in predicted estrogen-responsive regions is *WNT4*, a gene required for female development in other vertebrates. *WNT4* suppresses *SOX9* and 5-alpha reductase activity and promotes the formation of Müllerian ducts via frizzled receptor binding (Hsieh et al. 2002). Frizzled receptor genes *FZD2, FZD3, FZD6, FZD8*, and *FZD9* are all significantly expressed in the post-TSP gonads in both males and females. We therefore hypothesize that *WNT4* plays a role in sex differentiation in the American alligator similar to its role in other vertebrates, although unlike in vertebrates with genetic sex determination, its expression is determined by incubation temperature via estrogen signaling.

### Comparative assembly

We used the American alligator genome to scaffold the previously-published genome assemblies of two other crocodilians, the saltwater crocodile *Crocodylus porosus* and the gharial *Gavialis gangeticus*, based on synteny. These published assemblies have scaffold N50s of 205kb and 127kb, respectively. We performed comparative assembly on these genomes with Ragout (Kolmogorov 2013). Through this process, we were able to increase the scaffold N50 of the saltwater crocodile genome assembly from 205 kb to 84 Mb and the gharial genome assembly from 128 kb to 96 Mb. For comparison, the mean chromosome sequence length of the saltwater crocodile and gharial genomes are 117 Mb and 165 Mb, respectively.

To assess the accuracy of the synteny based scaffolding, we tested a random set of the scaffold joins predicted by Ragout for each species. We verified predicted scaffold joins using PCR with primers chosen such that the amplified regions would be unique in the genome assembly and would span the joins made by Ragout. We successfully amplified these gap regions for 18 out of 20 predicted joins tested in the saltwater crocodile genome and 22 out of 29 predicted joins tested in the gharial genome. Full results and primers used for join verification are in Table S5.

### Transposable elements

Repetitive sequences comprise more than one-third of the alligator genome assembly (Table S3). Almost a quarter of the genome is derived from just three TE superfamilies: LINE CR1s (12.2%) and the DNA transposons Harbinger (7.5%) and hAT (8.2%). TEs in general appear to accumulate more slowly in crocodilians than in other vertebrate taxa (excluding Testudines) and few new TE families, or even insertions, appear in any lineage of crocodilians since their divergence (Green et al. 2014; Suh et al. 2015). Data from the updated alligator assembly does not contradict these findings. Repeat content in general and from each of the dominant superfamilies are similar not only between alligator assemblies, but among crocodilians (Table S4), as determined by pre-masked genomes (http://repeatmasker.org/genomicDatasets/RMGenomicDatasets.html; accessed 15 March 2016). Only CR1 content varies between alligator assemblies to an appreciable degree. This variation, though, may be greater than it seems when contrasted with the near uniformity in the TE annotations across existing crocodilian assemblies (Green et al. 2014) (Table S4). Highly repetitive, nearly identical sequences are difficult to assemble from short reads and are likely underrepresented in genome assemblies, so an improved assembly may be able to identify these to a greater degree. Repeats in both alligator assemblies are biased towards those more than 10% diverged from their respective consensus element (Figure S5). No clear “burst” of CR1 activity specific to any one divergence bin is apparent, so it is likely that the additional CR1 insertions are distributed among elements with high and low mutation loads. Some of the variation in CR1 annotations between alligator assemblies is almost certainly due to stochasticity introduced by homology-based identification. Further it is possible that comparable improvements to the gharial and crocodile assemblies would yield similar changes in CR1 annotation.

### Small RNAs

MicroRNAs have been identified *de novo* in model vertebrate species, but for non-model species, miRNAs are usually identified based on sequence conservation with known miRNAs in other species. We sequenced a library of small RNAs isolated from alligator testis and used the resulting reads to predict 60 putative miRNAs after filtering for quality, including one, aca-mir-425, which appears in the American alligator, saltwater crocodile, and gharial genomes, but not in the chicken genome. See Supplemental Results for more details.

## Discussion

We present an improved assembly of the American alligator (*Alligator mississippiensis*) genome. After demonstrating its accuracy, we used this genome to examine synteny between the American alligator and chicken (*Gallus gallus*) genomes, improve the genomes of two other crocodilian species, and predict genomic regions likely to be under estrogenic regulation of gene expression in estrogen-responsive tissues. Finally, we showed that genes in these predicted estrogen-responsive regions are significantly more likely to have female-biased expression in post-TSP gonads. We thus conclude that the genomic architecture of estrogen signaling is remarkably well-conserved within vertebrates and that it is a fundamental early driver of female-biased gene expression in the post-TSP embryonic gonads of the American alligator.

The American alligator is an important study organism due to its ecological and economic importance and its temperature-dependent sex determination system which makes it especially susceptible to environmental contamination during development. The mechanisms responsible for temperature-dependent sex determination are not well-understood in any of the numerous species with this mode of sex determination.

Expression of aromatase, the enzyme that produces estrogen, has been hypothesized to be a master regulator of sex-biased gene expression in developing alligator embryos (Lance 2009). This is due to the ability of estrogen exposure to cause sex reversal in embryos incubated at male-producing temperature (Bull et al. 1988) and its extreme sex-biased expression in embryonic gonads after TSP (Gabriel et al. 2001). While much work is currently being performed to determine the pathway that allows aromatase expression to vary with temperature (Parrott et al. 2014; Yatsu et al. 2015; McCoy et al. 2016), less attention has been paid to the question of which genes estrogen regulates during sexual development in the American alligator or how estrogen regulates them despite its pivotal role early in embryonic sexual differentiation in alligator.

In this paper, we hypothesized that estrogen regulates gene expression in developing American alligator embryos through the same mechanism by which it is known to do so in humans, and that this mechanism can explain much of the female-biased gene expression that occurs after the temperature-sensitive period. Using the latest model of estrogen regulation of gene expression and *CTCF*-mediated chromatin looping in humans, we predicted the regions of the American alligator genome that are most likely to be under estrogenic regulation of gene expression. We then determined which genes have sex-biased expression in three tissues of developing alligator embryos at three different developmental time-points using RNA-sequencing to measure gene expression. We found strong support for our hypothesis based on the fact that genes contained in our predicted estrogen-responsive genomic regions are significantly more likely to show female-biased expression in the post-TSP gonads. Our results provide new evidence for Lance’s hypothesis that aromatase and its production of estrogen are a major driver of sex-biased gene expression in TSD in the American alligator (Lance 2009). Finally, we show that despite the different roles of estrogenic regulation of gene expression in sexual development between humans and alligators, much of the underlying mechanism responsible for estrogen regulation of gene expression is conserved between these two species.

Although our study does not fully elucidate the downstream effects of female-biased gene expression caused by estrogen signaling in the post-TSP gonads, *WNT4*’s female-biased expression and presence in a predicted estrogen-sensitive region provide a possible explanation for some of these effects. In mammals, *WNT4* expression prevents the formation of male-specific vasculature by preventing migration of endothelial and steroidogenic cells from mesonephros tissues to gonads (Jeays-Ward et al. 2003). It performs this action through upregulation of follistatin (Yao et al. 2004). *FST*, the gene coding for follistatin, is among the genes we find to have female-biased expression in the post-TSP alligator gonad, suggesting that *FST* may be among the genes indirectly regulated by estrogen signaling after the temperature-sensitive period. Furthermore, *WNT4* promotes expression of aromatase in mammals (Boyer et al. 2010). If the same is true in post-TSP embryonic alligator gonads, *WNT4* and aromatase may cooperate through a feed-forward mechanism in which estrogen promotes the expression of *WNT4* and *WNT4* promotes the expression of aromatase, which then creates more estrogen.

To fully understand the role of estrogenic regulation of gene expression in temperature-dependent sex determination in the American alligator, it will be necessary to go beyond *in silico* predictions of estrogen-responsive genomic regions and genes and verify our predictions *in vivo*. The human-based models we used are known to have far from perfect specificity even in humans. For example, a non-negligible number of experimentally determined ESR1 binding sites in the human genome do not match the canonical binding motif (Gruber et al. 2004; Carroll et al. 2006). We believe that our results provide sufficient support for our hypothesis to make such experiments worthwhile.

## Methods

### Sequencing and assembly

DNA was extracted with Qiagen blood and cell midi kits according to the manufacturer’s instructions. Briefly, cells were lysed and centrifuged to isolate the nuclei. The nuclei were further digested with a combination of Proteinase K and RNase A. The DNA was bound to a Qiagen genomic column, washed, eluted and precipitated in isopropanol, and pelleted by centrifugation. After drying, the pellet was resuspended in 200μL TE (Qiagen).

We generated the Chicago library as previously described by Putnam et al. (2016). Briefly, high-molecular weight DNA was assembled into chromatin *in vitro*, and then chemically cross-linked before being restriction digested. The overhangs were filled in with a biotinylated nucleotide, and the chromatin was incubated in a proximity-ligation reaction. The cross-links were then reversed, and the DNA purified from the chromatin. The library was then sonicated, and finished using the NEB Ultra library preparation kit (NEB cat # E7370), according to the manufacturer’s instructions, with the exception of a streptavidin bead capture step prior to indexing PCR. We sequenced the Chicago library on a single lane on the Illumina HiSeq 2500, resulting in 210 million read pairs.

The contig assembly was made with MERACULOUS (Chapman et al. 2011) and scaffolded using the Chicago library with Dovetail Genomics’ HiRise scaffolder as previously described by Putnam et al. (2016).

### Annotation

We made gene predictions using AUGUSTUS version 3.0.3 (Stanke et al. 2006). We provided as extrinsic evidence to AUGUSTUS RNA-seq alignments made using TopHat2 version 2.0.14 (Kim et al. 2013), repetitive region predictions made using RepeatScout (Price et al. 2005) and RepeatMasker Open-4.0 (Smit et al. 2015), and alignments of published chicken protein sequences made using Exonerate version 2.2.0 (Slater and Birney 2005). We assigned names to these predicted proteins and genes using reciprocal best hits BLAST searches between the set of predicted protein sequences and published protein sequences from related organisms. We also assigned Gene Ontology terms to our predicted proteins using InterProScan (Jones et al. 2014).

To annotate the genome for microRNAs, we extracted and purified small RNAs from testis tissue of a reproductively-mature alligator caught in the Rockefeller Wildlife Refuge (Grand Chenier, LA) using TRIzol reagent followed by an ethanol precipitation. We sequenced the resulting library on a MiSeq, and then, after filtering, used the miRDeep2 pipeline (Friedländer et al. 2012) and MapMi (Guerra-Assunção and Enright 2010) to align these sequences to and predict miRNAs in the alligator genome.

For more detail on our annotation process, see the Supplemental Methods.

### Egg harvesting, incubation, and dissection

All field and laboratory work were conducted under permits from the Florida Fish and Wildlife Conservation Commission and US Fish and Wildlife Service (Permit #: SPGS-1 0-44). 5 clutches of alligator eggs were collected from the Lake Woodruff National Wildlife Refuge, where relatively low chemical contamination of persistent organic pollutants allow American alligators to exhibit healthy reproductive activity. One egg from each clutch was dissected to identify the developmental stage of the embryo based on criteria described by Ferguson (1985). Eggs were incubated at 30°C (female-producing temperature, FPT) until they reached stage-19 based on an equation predicting their development (Kohno and Guillette 2013). At the predicted stage 19, which was before the temperature-sensitive period (stage 21-24) for alligator TSD (Lang and Andrews 1994), the incubation temperature was either kept constant at FPT or increased to 33°C (male-producing temperature, MPT). The alligator embryos were dissected and the gonad-adrenal-mesonephros (GAM) complex was isolated and preserved in ice-cold RNAlater (Ambion/Thermo Fisher Scientific) at 3 or 30 days after the stage 19. Gonadal tissues were carefully isolated from GAM under a dissection microscope after RNA stabilization in RNAlater, and stored at −80°C until RNA isolation.

### RNA-sequencing, expression quantification, and differential expression analysis

Total RNAs were then extracted from the GAM samples using TRIreagent LS (Sigma). Poly-A+ RNA sequencing libraries were made from each sample using the TruSeq RNA library preparation kit v1 (Illumina). A total of 60 libraries were created by PCR amplification with Illumina barcoding primers at 17 reaction cycles and quantified using a Bioanalyzer DNA 1000 kit (Agilent). Libraries were then pooled and sequenced on a HiSeq 2000 Sequencing system (Illumina).

We removed adapters from the reads using SeqPrep (https://github.com/jstjohn/SeqPrep) with default parameters and aligned them to the alligator genome using tophat2 (Kim et al. 2013) with default parameters. We used the cufflinks suite of tools to analyze gene expression (Trapnell et al. 2013). First, we quantified expression levels for each library using cuffquant with default parameters. None of the samples was developed enough to sex by sight, so to verify the sex of each sample, we looked at the relative expression of several markers commonly used to sex embryos (see Supplementary Methods and Figure S1). Finally, we used cuffdiff with an E-value cutoff of 0.01 to determine which genes were differentially expressed between males and females in each of the three tissues at both of the post-incubation time points.

### Predicting estrogen-responsive regions of the alligator genome

The DNA-binding domain of estrogen receptor alpha (*ESR1*) and the zinc fingers of *CTCF* are identically conserved in protein sequence among human, chicken, and alligator (Figure S3), suggesting that the DNA-binding motifs of these proteins are also conserved among these species. We predicted binding locations for these proteins by searching the alligator genome for sequences matching the human *ESR1*-binding motif (Lin et al. 2007) and the chicken CTCF-binding motif (Martin et al. 2011) using PoSSuM-search (Beckstette et al. 2006) with p-value cutoffs of 4.388 × 10^−6^ for *ESR1* and 1.214 × 10^−6^ for *CTCF*.

We considered any genomic region between two inward-facing CTCF motifs within 700kb to be possibly estrogen-responsive if it contained one or more ER-binding motifs.

### Synteny

We created synteny maps and calculated synteny statistics using SyMAP 4.2 (Soderlund et al. 2011), considering only scaffolds of at least 100kb and ordering the alligator scaffolds based on the chicken genome. We determined synteny for Galgal4 against both the previous version of the alligator genome (Green et al. 2014) and the updated alligator genome for comparison.

To calculate conservation of ordered gene n-lets between the alligator and chicken genomes, and the human and mouse genomes, we first found homologs in the second genome for genes in the first genome by performing a blastp search of the protein sequence of the primary isoform of each gene in the first genome against a database of all protein sequences in the second genome. We consider n-lets only of directly adjacent genes on the same scaffold. We then counted the counted the number of ordered gene n-lets in the first genome whose homologs also appear contiguously in the same order in the second genome.

### Comparative assembly

Nucleotide-level alignments between diverged genomes contain millions of small variations that are hard to analyse. To mitigate this, karyotype-level rearrangement studies typically use lower resolution alignments. Such alignments are described as a set of coarse “synteny blocks,” each such synteny block being a set of strand-oriented chromosome intervals in the set of genomes being compared that represents the homology relationship between large segments of the genomes. In this study, we also use synteny blocks to separate large structural variations from small polymorphisms. However, we take a hierarchical approach, with multiple sets of synteny blocks, each defined at a different resolution, from the coarsest, karyotype level all the way down to the fine grained base level. To create the hierarchy we use the principles developed by Sibelia tool (Minkin et al. 2013), which can create such a hierarchy for bacterial genomes, but adapted to use a multi-size A-Bruijn graph algorithm for constructing synteny blocks from a multiple genome alignment file in HAL format (Hickey et al. 2013), produced by Progressive Cactus (Paten et al. 2011).

At each level of resolution, the Ragout algorithm (Kolmogorov et al. 2014) decomposes the input genomes into a set of signed strings of synteny blocks, such that when the set of strings is concatenated together it forms a signed permutation of blocks. While each contiguous reference chromosome is transformed into a single sequence of signed synteny blocks, each chromosome in the target assembly corresponds to multiple sequences of synteny blocks because of the contig fragmentation. Due to this fragmentation, some information about the adjacencies between synteny blocks in the target genome is missing. Ragout constructs a breakpoint graph from the sets of synteny block strings and uses a rearrangement approach to infer these missing adjacencies, and also to detect misassembled contigs. Ragout further applies a “context-matching” algorithm to resolve repeats and puts them back into the previously constructed scaffolds. All these steps are performed iteratively starting from the lowest to the highest synteny block resolution. Scaffolds in the previous step are merged with scaffolds in the current step to refine the resulting scaffolds.

To assess the accuracy of joins, we designed primer pairs bracketing the gaps using Primer3 (Untergasser et al. 2012). We PCR amplified saltwater crocodile or gharial DNA with these primers at annealing temperatures ranging from 58°C to 62°C for 20 cycles. The joins, primers, and results are in Table S5.

### Transposable elements

We identified transposable elements and low complexity repetitive sequences in the alligator (A. *mississippiensis*) genome using RepeatMasker open-4.0 (Smit et al. 2015) and homology based searches with all known alligator repeats (RepBase Update v21.02). We created a repeat accumulation profile by calculating the Kimura 2-parameter (Kimura 1980) genetic distance between individual insertions and the homologous repeat in the *A. mississippiensis* library.

## Data Access

The new assembly of the American alligator genome is available on Genbank with the accession GCA_000281125.4. RNA-seq reads used to annotate protein-coding genes are available on SRA with the accession SRP057608. RNA-seq reads used for analysis of gene expression during sexual development are available on SRA under BioProject PRJNA322197. RNA-seq reads used to predict miRNAs are available on SRA under BioProject PRJNA285470.

## Acknowledgments

We thank the Florida Fish and Wildlife Conservation Commission and the US Fish and Wildlife Service for their assistance in obtaining collection permits; Steven Weber, Darrin Schultz, Stefany Rubio (UC Santa Cruz), and Jenny Korstian (Texas Tech University) for technical assistance; and Beth Shapiro (UC Santa Cruz) for discussion regarding the project.

